# Linking Diabetes mellitus to SARS-CoV-2 infection through differential targeting of the microRNAs in the Pancreas tissue

**DOI:** 10.1101/2021.03.31.437823

**Authors:** Bhavya, Ekta Pathak, Rajeev Mishra

## Abstract

Coronavirus Disease 2019 (COVID-19) severity and Diabetes mellitus affect each other bidirectionally. The plus-sense single-stranded RNA (+ssRNA) genome of the SARS-CoV-2 virus can be targeted and suppressed by the host cell’s microRNAs (miRNAs). Using the differential gene expression analysis between the mock-infected and the SARS-CoV-2-infected pancreatic tissue, we report five Diabetes-associated genes that are upregulated due to SARS-CoV-2 infection in the hESC pancreas tissues. Ten miRNAs regulating these five genes can potentially target the SARS-CoV-2 genome. We hypothesize that the SARS-CoV-2 genome copies in the infected human pancreas cell compete with the host cell’s native genes in being regulated by the native miRNAs. It leads to the reduced miRNA-regulation and, thus, the upregulation of the Diabetes-associated native genes. Thus, the resultant new-onset or elevated Diabetic symptoms may worsen the condition of COVID-19 patients.

## Introduction

Coronavirus Disease (COVID-19) is a fast-spreading disease that has caused a global crisis. This pandemic is a highly infectious viral disease caused by Severe Acute Respiratory Syndrome Coronavirus 2 (SARS-CoV-2), also known as novel coronavirus-2019 [1, 2]. According to the World Health Organization’s (WHO’s) COVID-19 Dashboard [3], 110,749,023 cases and 2,455,131 deaths have been reported globally due to COVID-19 as of 21^st^ February 2021. SARS-CoV-2 has a positive-sense single-stranded ribonucleic acid (+ssRNA) genome enclosed in a protein-containing lipid bilayer [4, 5]. Its RNA genome acts as a messenger RNA (mRNA) after entering the host cell and is directly translated by the host cell’s ribosomes, resulting in viral proteins. These translated viral proteins are also responsible for viral RNA replication [6].

Patients having either type 1 or type 2 Diabetes mellitus (commonly referred to as Diabetes) show poor prognosis with SARS-CoV-2 infection due to blood glucose level fluctuations and metabolic complications. An increase in blood glucose level helps in the SARS-CoV-2 replication and proliferation in the human monocytes [5, 7]. New-onset Diabetes has also been observed after COVID-19 infection [8]. Thus, COVID-19 severity and Diabetes affect each other in a bidirectional manner. Moreover, the resultant hyperglycaemia can diminish the individual’s immune response towards handling the viral infection. COVID-19 mortality is also amplified for diabetic patients due to complications like cardiovascular diseases and kidney-related problems [8-11]. Thus, regulating the blood glucose levels and preventing diabetic complications is required for diabetic COVID-19 patients to avert severe consequences of the infection on them [5].

Autopsy reports have been known to reveal COVID-19 infection in the human pancreas [12]. It becomes possible due to the angiotensin-converting enzyme 2 (ACE2) receptors on the pancreas cells. ACE2 acts as a very high-affinity receptor on the cell for the spike protein of SARS-CoV-2 [13, 14]. MicroRNAs (miRNAs) play an essential role in the regulation of gene expression in a cell. In a virus-infected cell, the host cell’s miRNAs help in cell defense by targeting the viral RNA genome or its transcribed RNAs. In humans, the miRNAs mostly target the 3’UTR of the mRNAs. However, the miRNAs primarily target the 3’UTR and 5’UTR of the viral RNA genomes. During a viral infection, the host cell’s miRNAs may target the viral genome rather than the native mRNAs. It leads to a competition for miRNA regulation between the host cell’s mRNAs and the viral genome copies in the cell. This differential targeting of the miRNAs may result in the dysregulation of the host cell’s genes [15-18].

In this study, we hypothesize the link between Diabetes and COVID-19 through differential targeting of the miRNAs in the Pancreas tissue. The disease enrichment based on differential gene expression analysis revealed pancreas- and diabetes-associated terms. We identified the miRNAs that can potentially target the SARS-CoV-2 genome as well as the Diabetes-associated genes upregulated in SARS-CoV-2-infected pancreas tissue.

## Materials and Methods

### Differential gene expression analysis

The RNA-Seq gene expression data for mock-infected and SARS-CoV-2-infected hESC pancreatic tissue were retrieved from NCBI. For the study, three mock-infected and three SARS-CoV-2-infected hESC pancreatic tissue samples were chosen from the Gene Expression Omnibus (GEO) dataset accession, GSE151803 [19]. The Differentially Expressed Genes (DEGs) between the mock-infected and the SARS-CoV-2-infected pancreatic tissue were obtained using the DESeq2 R package (padj < 0.05 and |log2FC| > 1) [20, 21].

### Disease enrichment analysis

DEGs-based disease enrichment analysis was done through DAVID – Functional Annotation Tool considering the Gene-Disease Associations Dataset (GAD) [22, 23]. The pancreas-associated diseases and the DEGs associated with them were selected for further studies. A literature search was done for the obtained DEGs to understand their role in the respective diseases.

### Gene-gene interaction network

The gene-gene interaction (GGI) network of the DEGs was created using the Cytoscape-GeneMANIA app [24-26]. Network topological analysis was done using the Cytoscape-NetworkAnalyzer plugin [27] to get an insight about the influence of the genes in the network.

### Differentially targeting miRNAs

The complete genome reference sequence of SARS-CoV-2, Wuhan-Hu-1, was retrieved from NCBI RefSeq ID, NC_045512.2 [28]. The 3’UTR and 5’UTR of the viral genome were obtained. The human miRNAs targeting the 3’ and 5’ UTR of the SARS-CoV-2 genome (CoV-tar-miRNAs) were obtained using the miRDB online tool [29, 30]. The target genes of CoV-tar-miRNAs were obtained using the Predicted Target Module of miRWalk 2.0 [31, 32].

### Diabetes-linked CoV-tar-miRNAs

The target genes of the CoV-tar-miRNAs were compared to the enriched Diabetes-associated genes. The genes common in both were those Diabetes-associated genes that may get upregulated in pancreas cells due to miRNAs’ differential targeting during SARS-CoV-2 infection. Furthermore, the availability of miRNAs in the pancreas tissue was checked from the web-based repository, Tissue-Atlas [33].

## Results

### Differential Gene Expression between mock-infected and SARS-CoV-2-infected pancreatic tissue

Using the DESeq2 R package, 30 differentially expressed genes (DEGs) between the mock-infected and SARS-CoV-2-infected pancreatic tissues of the GEO dataset GSE151803 were revealed [19-21]. 26 DEGs were upregulated and 4 DEGs were downregulated (Table 1) in the SARS-CoV-2-infected hESC pancreas tissue.

**Table 1:** Differentially Expressed Genes in SARS-CoV-2-infected hESC pancreas tissue.

| S. No. | Gene Symbol | Log2FoldChange | S. No. | Gene Symbol | Log2FoldChange |
| --- | --- | --- | --- | --- | --- |
| 1. | CP | 3.218646021 | 16. | CHRD | 1.226960717 |
| 2. | SOCS3 | 2.897737916 | 17. | GADD45G | 1.223936929 |
| 3. | VNN3 | 2.676974204 | 18. | APOL6 | 1.164596924 |
| 4. | CEBPD | 2.041519486 | 19. | SAMD11 | 1.129551911 |
| 5. | FAM169B | 1.887720767 | 20. | AGT | 1.106392771 |
| 6. | VNN2 | 1.650151224 | 21. | C9orf16 | 1.096117015 |
| 7. | HPX | 1.576405664 | 22. | NLRC5 | 1.094272111 |
| 8. | EVA1A | 1.483037767 | 23. | SUCNR1 | 1.067266654 |
| 9. | SERTM1 | 1.482242018 | 24. | PSMB8 | 1.031089013 |
| 10. | CXCL2 | 1.464664038 | 25. | UNC5CL | 1.015471709 |
| 11. | C8B | 1.45102627 | 26. | MAP6D1 | 1.000047502 |
| 12. | CFB | 1.449773368 | 27. | AKR1B10* | -1.942847572 |
| 13. | SERPINA3 | 1.282619509 | 28. | SYNDIG1L* | -1.628255609 |
| 14. | TACR1 | 1.253706268 | 29. | PKHD1L1* | -1.414217069 |
| 15. | HCP5 | 1.228387286 | 30. | TMEM236* | -1.040226661 |

\*Genes downregulated in SARS-CoV-2-infected hESC pancreas tissue.

### Disease enrichment of DEGs

DAVID Tool was used for DEGs-based disease enrichment analysis based on the Gene-Disease Associations Dataset (GAD) [22, 23]. Gene-based disease enrichment of the 30 DEGs resulted in Type I Diabetes as the most significant GAD disease. It was linked to four upregulated DEGs, i.e., CP, SOCS3, AGT and PSMB8 (Table 2). Two upregulated DEGs, CP and CFB were enriched for the term “insulin.” COVID-19 has been linked with the other resultant disease terms, but only the terms associated with the pancreas were selected for the study.

**Table 2:** Gene-based GAD disease enrichment of DEGs.

| S. No. | Disease Term | Genes |
| --- | --- | --- |
| 1. | **Type 1 Diabetes** | SOCS3, AGT<sup>S</sup>, CP<sup>S</sup>, PSMB8 |
| 2. | Alzheimer's Disease, Parkinson's Disease, **Insulin**, Lung Function, Depression, Longevity | CFB<sup>S</sup>, CP<sup>S</sup> |
| 3. | Cerebrovascular Disease | AGT<sup>S</sup>, SERPINA3<sup>S</sup> |
| 4. | Respiratory Syncytial Virus Bronchiolitis, Asthma, Bronchiolitis | SOCS3, CXCL2<sup>S</sup>, PSMB8 |
| 5. | Birth Weight, Leukemia, Acute Myeloid Leukemia, Precursor Cell Lymphoblastic Leukemia-Lymphoma, Meningeal Neoplasms, Meningioma, Non-Hodgkin's Lymphoma | C8B<sup>S</sup>, SOCS3, CFB<sup>S</sup> |
| 6. | Nephropathy | AGT<sup>S</sup>, AKR1B10<sup>S</sup> |
| 7. | Myocardial Infarction | EVA1A, AGT<sup>S</sup>, GADD45G, SERPINA3<sup>S</sup> |
| 8. | Plasma HDL Cholesterol (HDL-C) Levels | SOCS3, CEBPD, AGT<sup>S</sup> |
| 9. | Macular Degeneration | C8B<sup>S</sup>, CFB<sup>S</sup>, SERPINA3<sup>S</sup> |

<sup>S</sup> Genes encoding for secretory proteins.

### Gene-Gene interaction (GGI) network

The Cytoscape-GeneMANIA [24-26] GGI network of the DEGs contains 29 genes as HCP5 is a lncRNA, and its data is not present in GeneMANIA. Diabetes-associated (Type 1 Diabetes and insulin-associated) DEGs are connected directly or indirectly through co-expression (Figure 1, Table 3). Two downregulated DEGs, i.e., AKR1B10 and PKHD1L1, show interaction with the upregulated DEGs (Figure 1, Table 4).

**Table 3:**
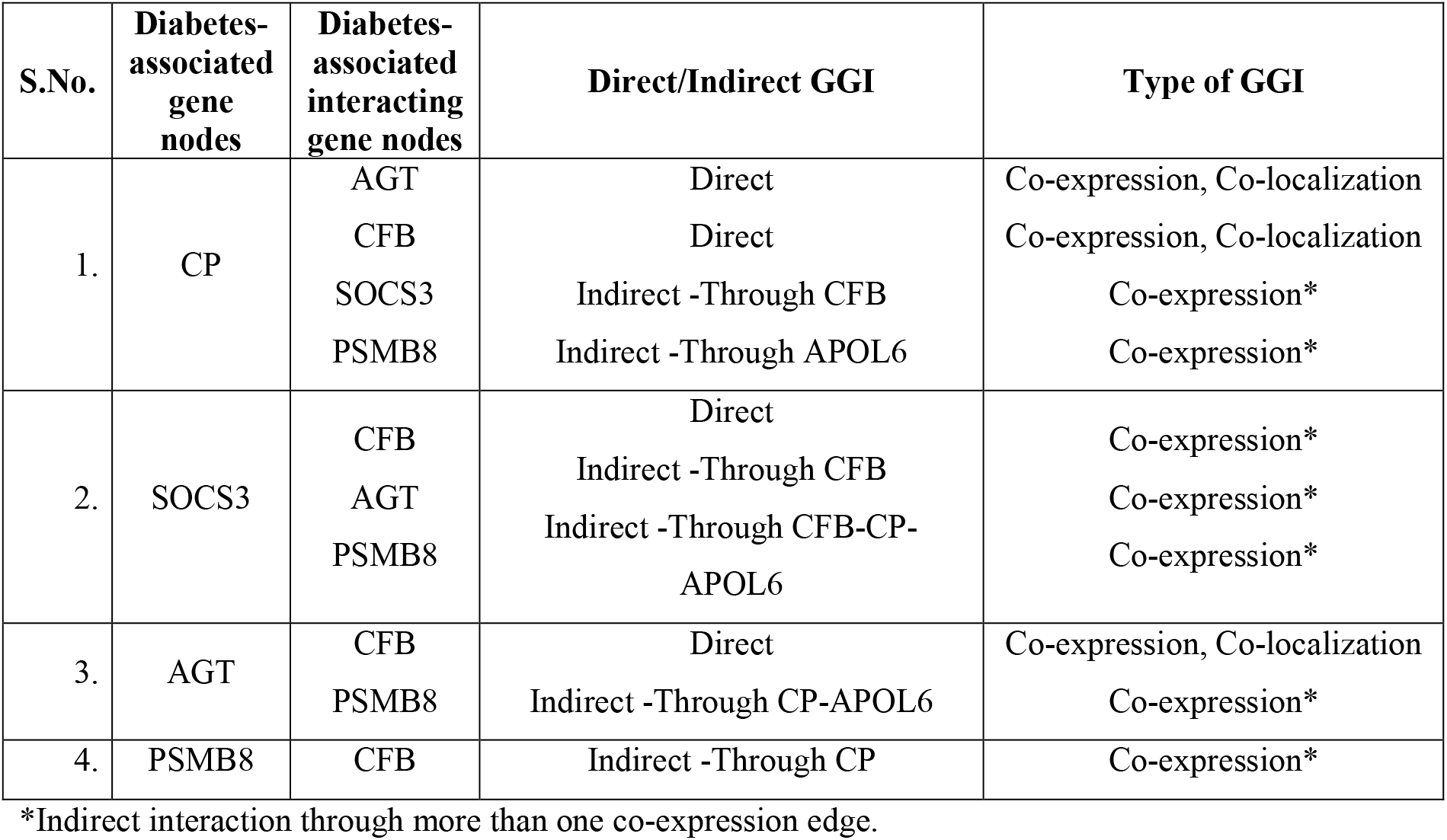
GeneMANIA GGI between the Diabetes-associated DEGs.

**Table 4:** GeneMANIA direct GGI between the upregulated and downregulated DEGs.

| S.No. | Downregulated gene | Upregulated gene | Direct/Indirect GGI | Type of GGI |
| --- | --- | --- | --- | --- |
| 1. | AKR1B10 | C8B | Direct | Co-expression |
| 2. | PKHD1L1 | CP | Direct | Shared protein domains |
| 3. | SYNDIG1L | None | - | - |
| 4. | TMEM236 | None | - | - |

**Figure 1.**
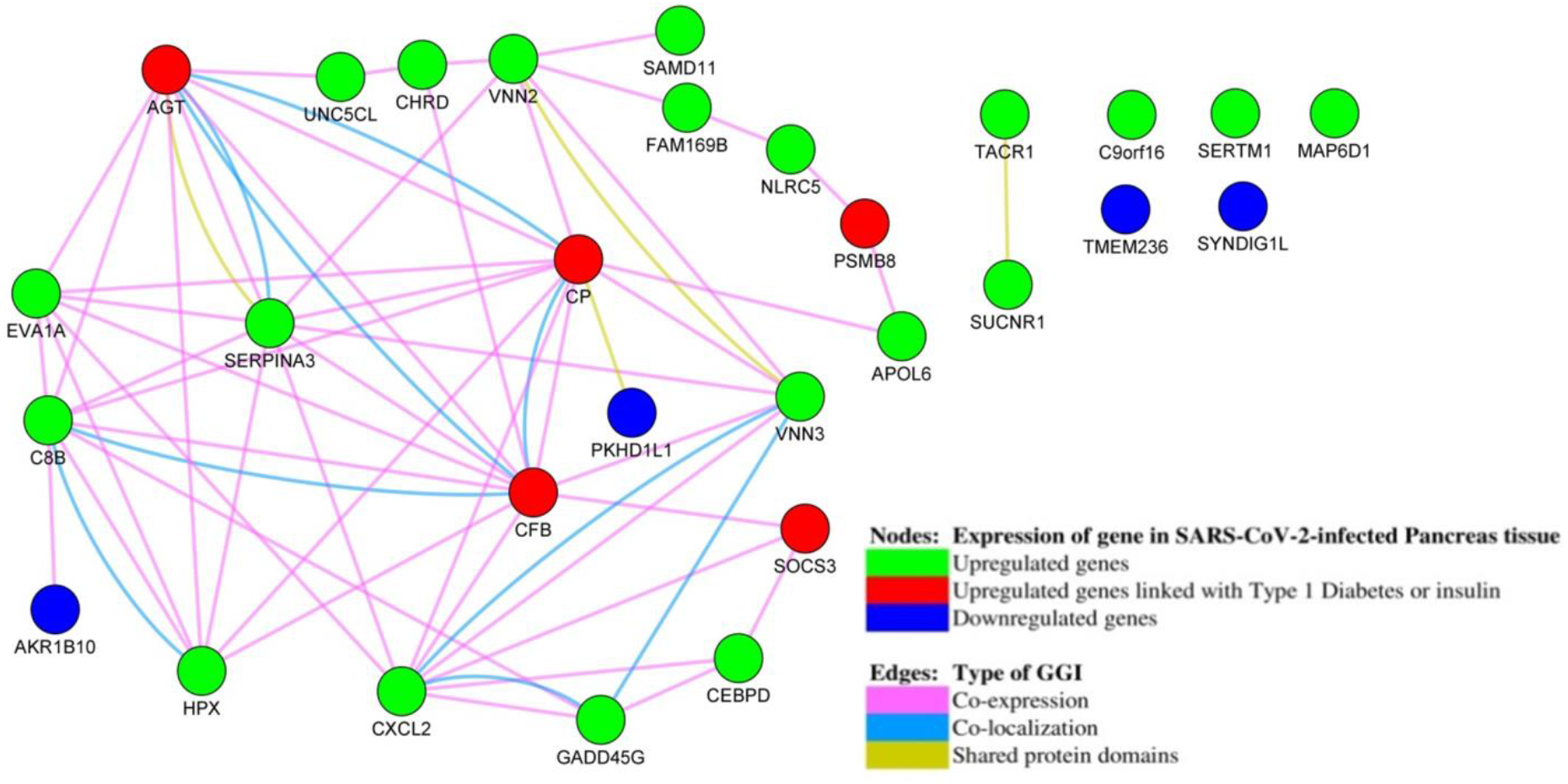
GeneMANIA gene-gene interaction network between the Differentially expressed genes.

After removing the duplicate edges between the gene nodes, the network’s topological parameters were calculated (Table 5). Closeness centralities of the SUCNR1 and TACR1 genes were ignored due to their interaction with each other only and not with the significant network. The Diabetes-associated gene CP has the highest closeness and betweenness centrality; thus, it is the most influential gene in the network. The CFB gene is the second-highest diabetes-associated gene in terms of closeness centrality.

**Table 5:** Topological parameters of the GGI network (in descending order of Closeness Centrality)

| S. No. | Gene name | Degree | Closeness Centrality | Betweenness Centrality |
| --- | --- | --- | --- | --- |
| 1. | SUCNR1 | 1 | 1 | 0 |
| 2. | TACR1 | 1 | 1 | 0 |
| 3. | CP<sup>D</sup> | 11 | 0.65625 | 0.334650416 |
| 4. | SERPINA3 | 9 | 0.6 | 0.078909675 |
| 5. | CFB<sup>D</sup> | 10 | 0.567567568 | 0.124459562 |
| 6. | C8B | 8 | 0.538461538 | 0.124801587 |
| 7. | EVA1A | 7 | 0.525 | 0.007420635 |
| 8. | VNN3 | 6 | 0.525 | 0.055616024 |
| 9. | VNN2 | 6 | 0.525 | 0.25037037 |
| 10. | AGT<sup>D</sup> | 7 | 0.512195122 | 0.062619048 |
| 11. | CXCL2 | 8 | 0.512195122 | 0.101135676 |
| 12. | HPX | 6 | 0.5 | 0 |
| 13. | CHRD | 3 | 0.446808511 | 0.034950869 |
| 14. | APOL6 | 2 | 0.4375 | 0.096216931 |
| 15. | GADD45G | 4 | 0.4375 | 0.022896825 |
| 16. | PKHD1L1 | 1 | 0.403846154 | 0 |
| 17. | SOCS3<sup>D</sup> | 3 | 0.396226415 | 0.007539683 |
| 18. | UNC5CL | 2 | 0.381818182 | 0.002380952 |
| 19. | FAM169B | 2 | 0.375 | 0.07521164 |
| 20. | CEBPD | 3 | 0.368421053 | 0.002380952 |
| 21. | AKR1B10 | 1 | 0.355932203 | 0 |
| 22. | SAMD11 | 1 | 0.35 | 0 |
| 23. | PSMB8<sup>D</sup> | 2 | 0.328125 | 0.023994709 |
| 24. | NLRC5 | 2 | 0.291666667 | 0.013492063 |
| 25. | C9orf16* | 0 | 0 | 0 |
| 26. | TMEM236* | 0 | 0 | 0 |
| 27. | SYNDIG1L* | 0 | 0 | 0 |
| 28. | SERTM1* | 0 | 0 | 0 |
| 29. | MAP6D1* | 0 | 0 | 0 |

\*Single node in the network.
<sup>D</sup> Differentially Expressed Genes associated with Diabetes.

### 3’UTR and 5’UTR of the viral genome

The SARS-CoV-2 genome consists of a linear 29903 nucleotides long ss-RNA [28]. The 3’UTR of the genome is 229 nucleotides long, including a polyA tail. It lies in the viral genome at the position from 29675 to 29903 nucleotides. The 5’UTR of the genome is 265 nucleotides long and spans from 1 to 265 nucleotides in the viral genome. Both 3’UTR and 5’UTR of the SARS-CoV-2 genome sequence is appended below:

>NC_045512.2:29675-29903 Severe acute respiratory syndrome coronavirus 2 isolate Wuhan-Hu-1, complete genome

CAATCTTTAATCAGTGTGTAACATTAGGGAGGACTTGAAAGAGCCACCACATTTTCACCGAGGCCACGCG GAGTACGATCGAGTGTACAGTGAACAATGCTAGGGAGAGCTGCCTATATGGAAGAGCCCTAATGTGTAAA ATTAATTTTAGTAGTGCTATCCCCATGTGATTTTAATAGCTTCTTAGGAGAATGACAAAAAAAAAAAAAA AAAAAAAAAAAAAAAAAAA

>NC_045512.2:1-265 Severe acute respiratory syndrome coronavirus 2 isolate Wuhan-Hu-1, complete genome

ATTAAAGGTTTATACCTTCCCAGGTAACAAACCAACCAACTTTCGATCTCTTGTAGATCTGTTCTCTAAA CGAACTTTAAAATCTGTGTGGCTGTCACTCGGCTGCATGCTTAGTGCACTCACGCAGTATAATTAATAAC TAATTACTGTCGTTGACAGGACACGAGTAACTCGTCTATCTTCTGCAGGCTGCTTACGGTTTCGTCCGTG TTGCAGCCGATCATCAGCACATCTAGGTTTCGTCCGGGTGTGACCGAAAGGTAAG

### miRNAs targeting the UTRs of the viral genome

Using miRWalk 2.0 online tool [31, 32], 10 and 11 miRNAs, i.e., CoV-tar-miRNAs, were found to be potentially targeting the 3’UTR (Table 6) and 5’UTR (Table 7) of the SARS-CoV-2 genome, respectively.

**Table 6:**
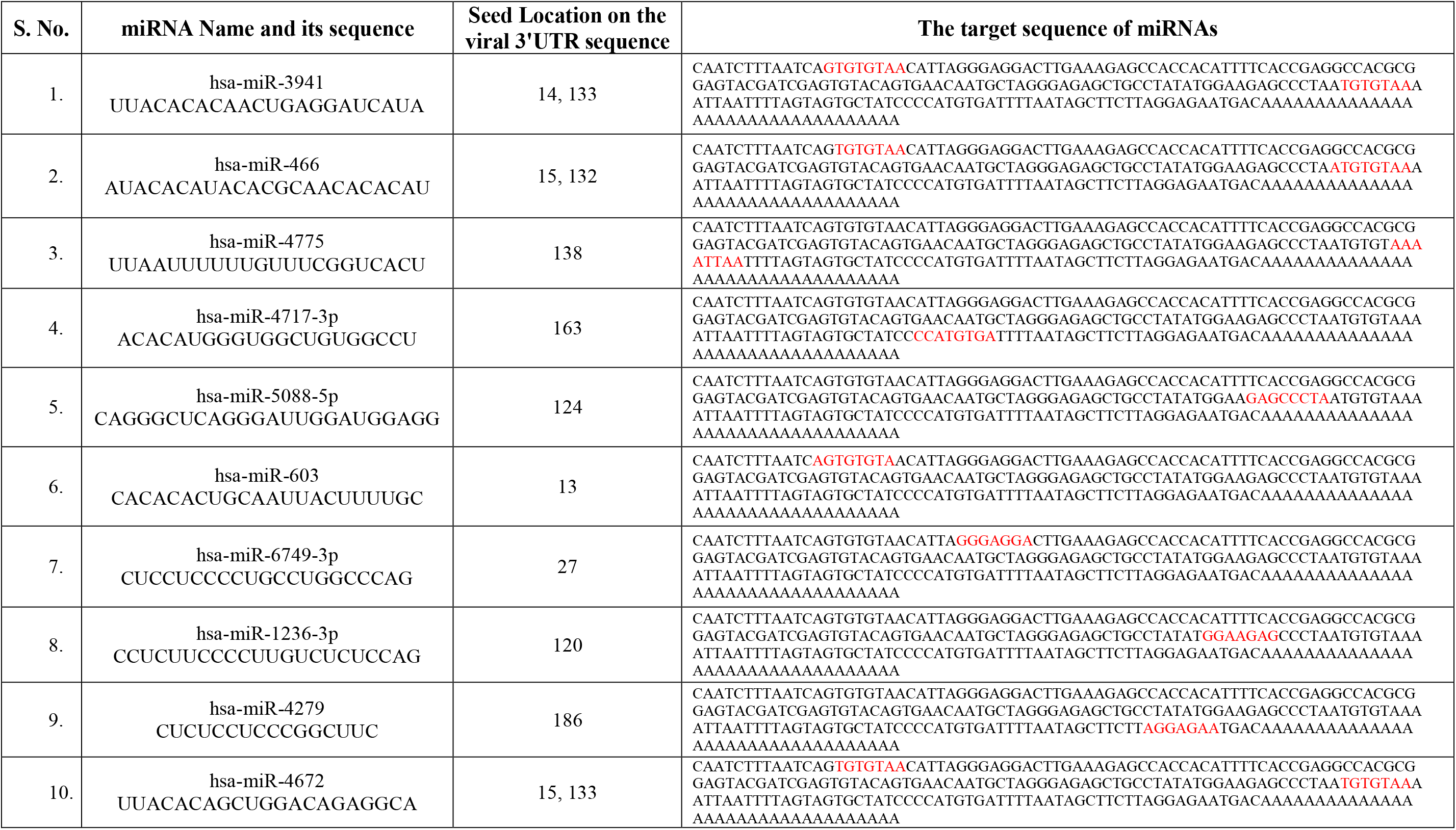
Potential miRNAs targeting the 3’UTR of the viral genome (CoV-tar-miRNAs)

**Table 7:**
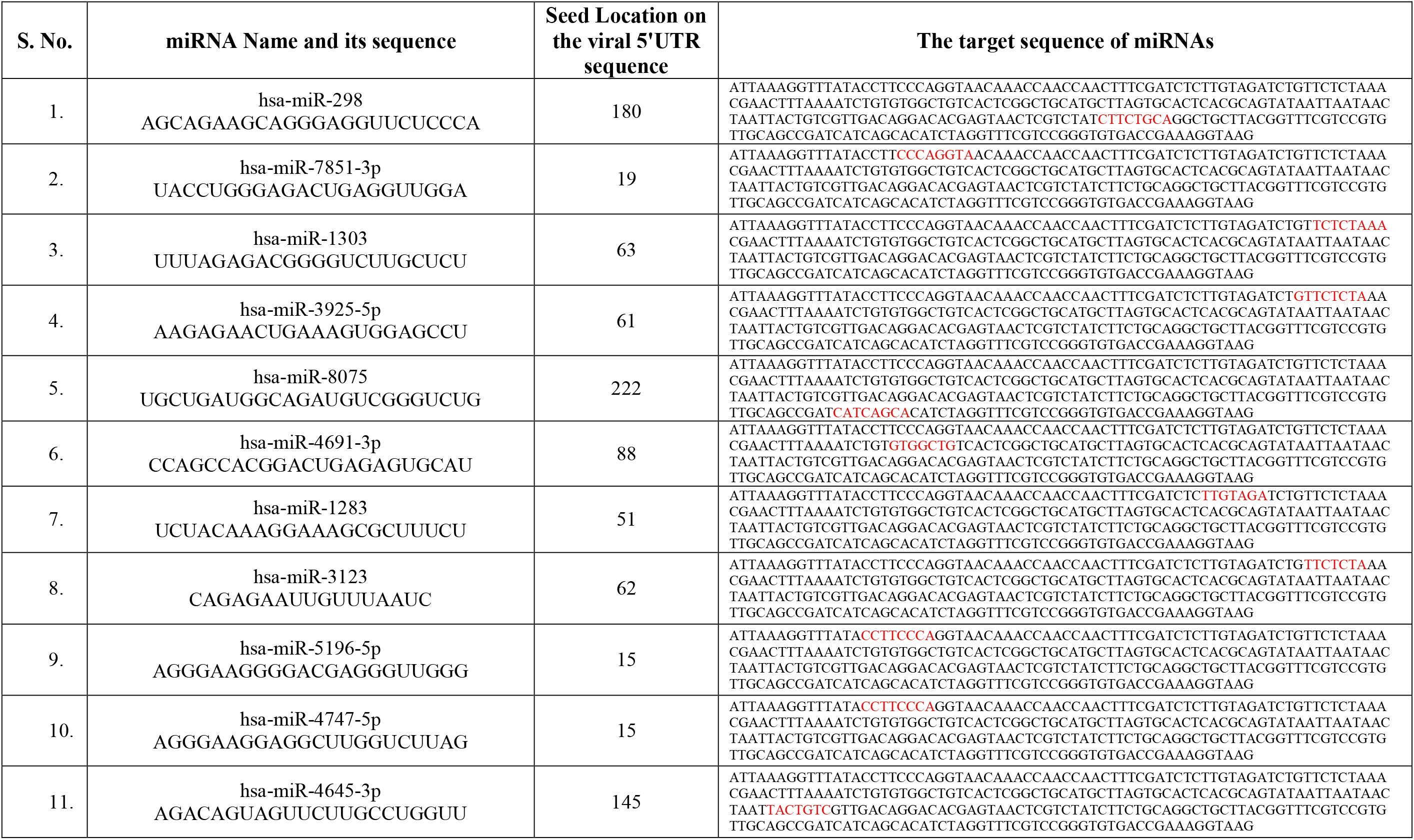
Potential miRNAs targeting the 5’UTR of the viral genome (CoV-tar-miRNAs)

### CoV-tar-miRNAs targeting Diabetes-associated DEGs

Each of the five Diabetes-associated DEGs was found to be targeted by one or more CoV-tar-miRNAs. Eight 3’UTR and nine 5’UTR CoV-tar-miRNAs were found to be potentially targeting the Diabetes-associated DEGs, respectively. Availability and raw microarray expression of five 3’UTR and six 5’UTR CoV-tar-miRNAs in the human pancreas were obtained from the TissueAtlas web-based repository. Due to the microarray expression data’s unavailability in TissueAtlas, hsa-miR-466, hsa-miR-3123, hsa-miR-4691-3p, hsa-miR-3941, hsa-miR-6749-3p and hsa-miR-7851-3p were not considered for further study. hsa-mir-4775 has a negative raw microarray expression value in the human pancreas; thus, it was also not considered for differential targeting analysis. Four 3’UTR and six 5’UTR CoV-tar-miRNAs are expressed in the human pancreas (Table 8).

**Table 8:** CoV-tar-miRNAs targeting the DEGs linked to Diabetes:

| S. No. | Diabetes-associated DEGs | SARS-CoV-2genome's position targeted by CoV-tar-miRNAs | CoV-tar-miRNAs targeting the DEGs | CoV-tar-miRNAs' raw microarray expression in human Pancreas |
| --- | --- | --- | --- | --- |
| 1. | CP | 3'UTR | hsa-miR-466*<br>hsa-miR-4775** | Unknown<br>-0.00390 |
|  |  | 5'UTR | hsa-miR-3123*<br>hsa-miR-4691-3p*<br>hsa-miR-4747-5p<br>hsa-miR-5196-5p | Unknown<br>Unknown<br>10.20753<br>80.05370 |
| 2. | SOCS3 | 3'UTR | hsa-miR-1236-3p<br>hsa-miR-3941*<br>hsa-miR-4279<br>hsa-miR-4717-3p<br>hsa-miR-5088-5p<br>hsa-miR-6749-3p* | 6.40509<br>Unknown<br>3.16020<br>20.06552<br>68.77350<br>Unknown |
|  |  | 5'UTR | hsa-miR-1303<br>hsa-miR-3123*<br>hsa-miR-3925-5p<br>hsa-miR-4691-3p*<br>hsa-miR-7851-3p* | 5.12508<br>Unknown<br>27.36460<br>Unknown<br>Unknown |
| 3. | AGT | 3'UTR | hsa-miR-4775* | -0.00390 |
|  |  | 5'UTR | hsa-miR-1303<br>hsa-miR-298<br>hsa-miR-4645-3p<br>hsa-miR-7851-3p* | 5.12508<br>0.88648<br>1.34874<br>Unknown |
| 4. | PSMB8 | 3'UTR | hsa-miR-1236-3p<br>hsa-miR-4279<br>hsa-miR-5088-5p | 6.40509<br>3.16020<br>68.77350 |
|  |  | 5'UTR | hsa-miR-298<br>hsa-miR-3123*<br>hsa-miR-3925-5p | 0.88648<br>Unknown<br>27.36460 |
| 5. | CFB | 3'UTR | hsa-miR-4775** | -0.00390 |

\* Microarray expression data not available on TissueAtlas
\*\* Not expressed in pancreas

## Discussion

The SARS-CoV-2 infection has been detected in many different human organs, including the pancreas. The viral infection may lead to dysregulation of genes in the infected cells. 26 and 4 genes were found to be upregulated and downregulated, respectively, in the SARS-CoV-2-infected hESC pancreas tissue. Among them, five upregulated genes, i.e., CP, SOCS3, AGT, PSMB8, and CFB, are associated with Diabetes (Type 1 Diabetes or insulin). Among the genes associated with other enriched diseases, six upregulated genes, i.e., CP, AGT, CFB, SERPINA3, CXCL2, and C8B, encode for proteins that are secreted to blood. These proteins may also reach other organs of the body, thus, being involved in other diseases. Their roles in other diseases must be further investigated. Diabetes can result in Nephropathy or Macular Degeneration [34-36]; therefore, they can be the indirect results of COVID-19.

The human miRNAs mostly target the 3’UTR of the mRNAs in the cytoplasm of the cell. However, they primarily target the 3’UTR and 5’UTR of the infecting viral RNA genome. It has been seen that the SARS-CoV-2 virus also infects the pancreas, among other organs of the human body. Since it is a +ss-RNA genome virus, its genome itself acts as an mRNA. Thus, the pancreas cell’s miRNAs must target the viral 3’UTR or 5’UTR, causing them to deflect from regulating the pancreas host cell’s native genes. This may cause the upregulation of the host cell’s native genes. Twenty-one human miRNAs (CoV-tar-miRNAs) were found to be potentially targeting the UTRs of the viral genome. Considering the availability of these miRNAs in the human pancreas tissue, 10 of them also target the Diabetes-associated genes, thus, keeping their expression in check before the infection. After infection, the SARS-CoV-2 genome competes with the mRNAs of these genes in being targeted and regulated by the miRNAs. This differential targeting of the miRNAs explains the upregulation of the Diabetes-associated genes after the viral infection (Figure 2).

**Figure 2.**
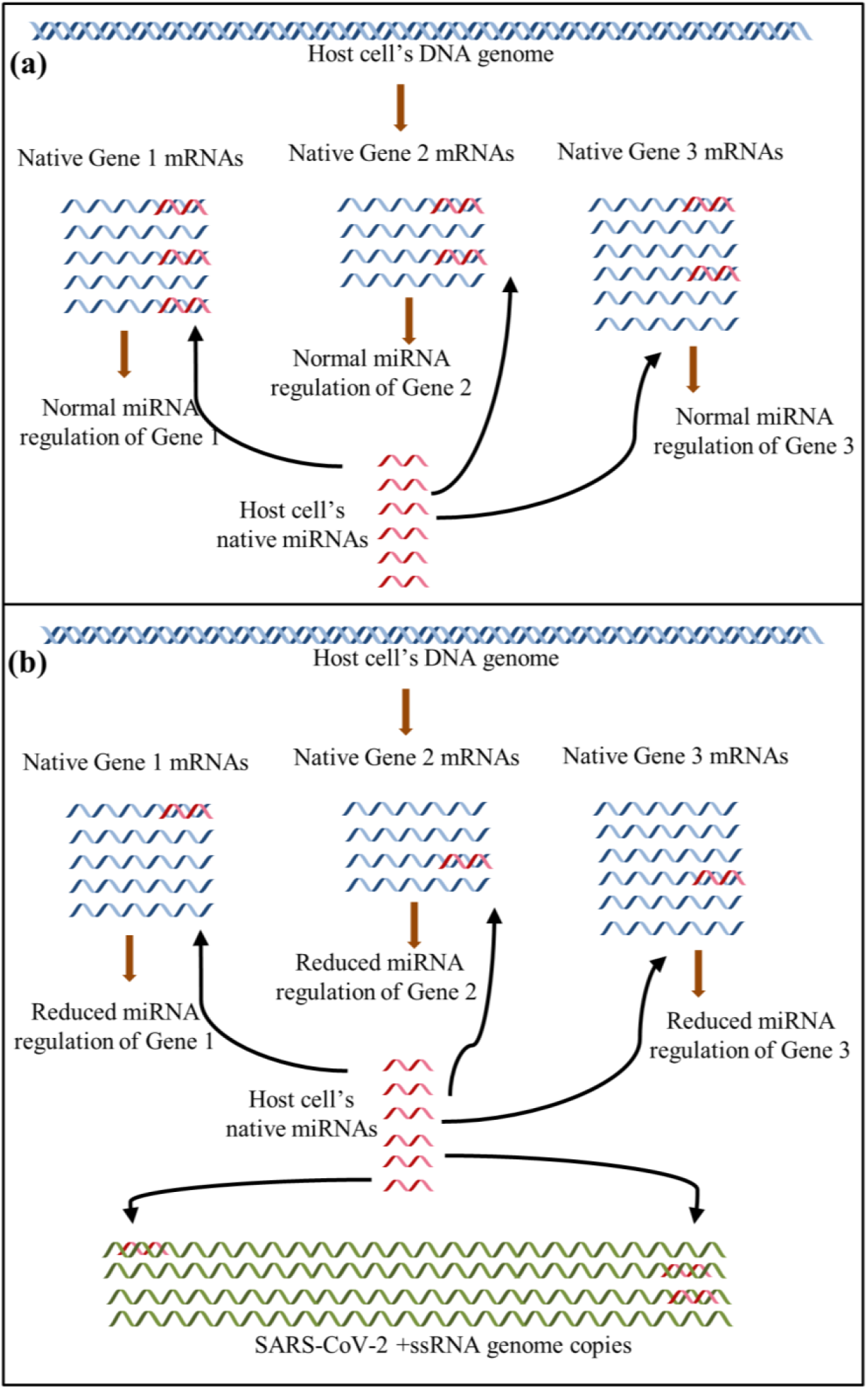
Differential targeting of the host cell’s native miRNAs. (a) The cell’s native miRNAs regulate the genes by targeting and suppressing the cell’s native mRNAs only. (b) In a SARS-CoV-2-infected cell, the viral +ssRNA genome copies compete with the host cell’s native mRNAs in being targeted by the native miRNAs. The native miRNAs get apportioned in targeting the SARS-CoV-2 genome copies too. It reduces the contribution of miRNAs in regulating the native cell’s genes, thus, upregulating them.

The SOCS3 (Suppressor of Cytokine Signaling 3) gene codes for a protein that helps regulate cytokine signal transduction [37]. It is the potential target of four 3’UTR CoV-tar-miRNAs, i.e., hsa-miR-1236-3p, hsa-miR-4279, hsa-miR-4717-3p and hsa-miR-5088-5p, and two 5’UTR CoV-tar-miRNAs, i.e., hsa-miR-1303 and hsa-miR-3925-5p. Thus, SOCS3 upregulation after the viral infection may be due to the differential targeting of these miRNAs. Overexpression of SOCS3 has been observed in mice having Type 1 Diabetes. SOCS3-deficiency in pancreas beta cells is associated with increased resistance to apoptosis, thus, preventing Type 1 Diabetes [38]. However, in the SARS-CoV-2-infected Pancreas tissue, the expression of SOCS3 may be increased due to differential targeting of miRNAs; thus, it decreases the beta cells’ resistance to the development of Diabetes.

The PSMB8 gene is the potential target of three 3’UTR CoV-tar-miRNAs, i.e., hsa-miR-1236-3p, hsa-miR-4279 and hsa-miR-5088-5p, and two 5’UTR CoV-tar-miRNAs, i.e., hsa-miR-3925-5p and hsa-miR-298. PSMB8 codes for Proteasome 20S Subunit Beta 8 and has been found to promote apoptosis [39, 40]. Its upregulation may also contribute to the apoptosis of the pancreas’ beta cells.

The CP gene is the potential target of two 5’UTR CoV-tar-miRNAs, i.e., hsa-miR-4747-5p and hsa-miR-5196-5p. The CP gene encodes a secretory plasma protein called Ceruloplasmin, the level of which is found to be increased in the Diabetic condition [41]. CP has the highest closeness and betweenness centrality, indicating its strong influence on the network.

The CFB gene encodes for the secretory complement factor b, which links obesity to Diabetes. Its level is found to be increased during obesity. It is also linked to insulin resistance [42]. Although in our study, CFB gene expression does not seem to be affected by differential miRNAs targeting, its upregulation is linked with the upregulation of CP, SOCS3, and AGT genes as all these genes are co-expressed according to the gene-gene interaction network (Figure 1).

The AGT gene is the potential target of three 5’UTR CoV-tar-miRNAs, i.e., hsa-miR-1303, hsa-miR-298, and hsa-miR-4645-3p. AGT gene encodes for the secretory pre-angiotensinogen or angiotensinogen precursor, which is essential for the renin-angiotensin system to maintain the blood pressure and fluid and electrolyte homeostasis in the body [43, 44].

Our study suggests that after SARS-CoV-2 infection, these genes associated with Diabetes and cell death get upregulated due to differential miRNA targeting and lead the pancreas cells to death. This hypothesis can be applied to the insulin-producing pancreas beta cells, resulting in low or no insulin secretion. This can worsen the condition of COVID-19 patients due to diabetic complications or new onset of Diabetes, even if the pancreas cell’s miRNAs block the viral genome function. The mechanism of differential miRNA targeting must be further validated *in vitro*. For therapeutic purposes, artificial miRNAs can be designed and inserted into the infected cells to bind with the viral genome, thus, blocking both its function as well as the differential targeting of the host cell’s miRNAs. A research study with this concept has been done with four artificial miRNAs for targeting and blocking the Chikungunya virus genome [45].

## Conclusion

This study suggests that in a SARS-CoV-2-infected human pancreas cell, the native miRNAs target the viral genome instead of the cell’s mRNAs that were being targeted before the infection. This differential miRNA targeting causes the pancreas cell’s genes associated with Diabetes to upregulate, leading to diabetic complications or even new onset of Diabetes. Preventive, therapeutic methods are needed to block the viral genome from binding the host cell’s miRNAs and facilitate binding with the externally provided artificial miRNAs.

## Acknowledgement

B is supported with a Junior Research Fellowship from the Indian Council of Medical Research (ICMR), Govt. of India, New Delhi.

